# *Hic sunt dracones*: ancient DNA from the extinct palaeofauna of the tiny Mediterranean archipelago of Columbretes and its implications for human visitation

**DOI:** 10.64898/2026.10.30.763138

**Authors:** Gonzalo Oteo-García, Rafael Marquina-Blasco, Marta Calvo Revuelta, Sofia Selvatici, Isabelle Sofie Feinauer, Elisa Rondoni, Adrián Talavera, Sofia Panella, Ana Fagoaga, Francisco Javier Ruiz-Sánchez, David López Serrano, Gustau Aguilella, Anna Linderholm, Laura Parducci, Anders Götherström, Love Dalén

## Abstract

The Columbretes archipelago in the Mediterranean is an example of deep isolation and sporadic human interaction during the Holocene. We recovered ancient DNA from 2,600-year-old microfauna and historical museum specimens to investigate the island’s extinct fauna. Despite challenging preservation conditions, we successfully recovered mitogenomes from historical *Vipera latastei* specimens, and partial mitogenomes from snake, skink, and lizard subfossils. Phylogenetic analyses confirm the extinct Columbretes snake as *Vipera latastei*, sharing a haplotype with the historical specimen and clustering with modern populations from northeastern Iberia, validating its provenance and suggesting a single mitochondrial lineage since isolation. The extinct skink, *Chalcides bedriagai*, also shows affinities with eastern Iberia. These results point to a vicariant origin for the herpetofauna, likely isolated on the island following sea level rise after the Last Glacial Maximum. Conversely, a 2,600-year-old mouse incisor yielded a *Mus musculus* mitogenome, indicating Iron Age human-mediated introduction. Additionally, genomic analysis of a 11th-century CE human burial revealed an individual with genetic affinities to the Central Mediterranean, highlighting medieval maritime mobility. This study resolves the long-debated Columbretes snake identity, traces biogeographic origins and illustrates the interplay of natural isolation and anthropogenic influence on the island biodiversity.

**GRAPHICAL ABSTRACT:** 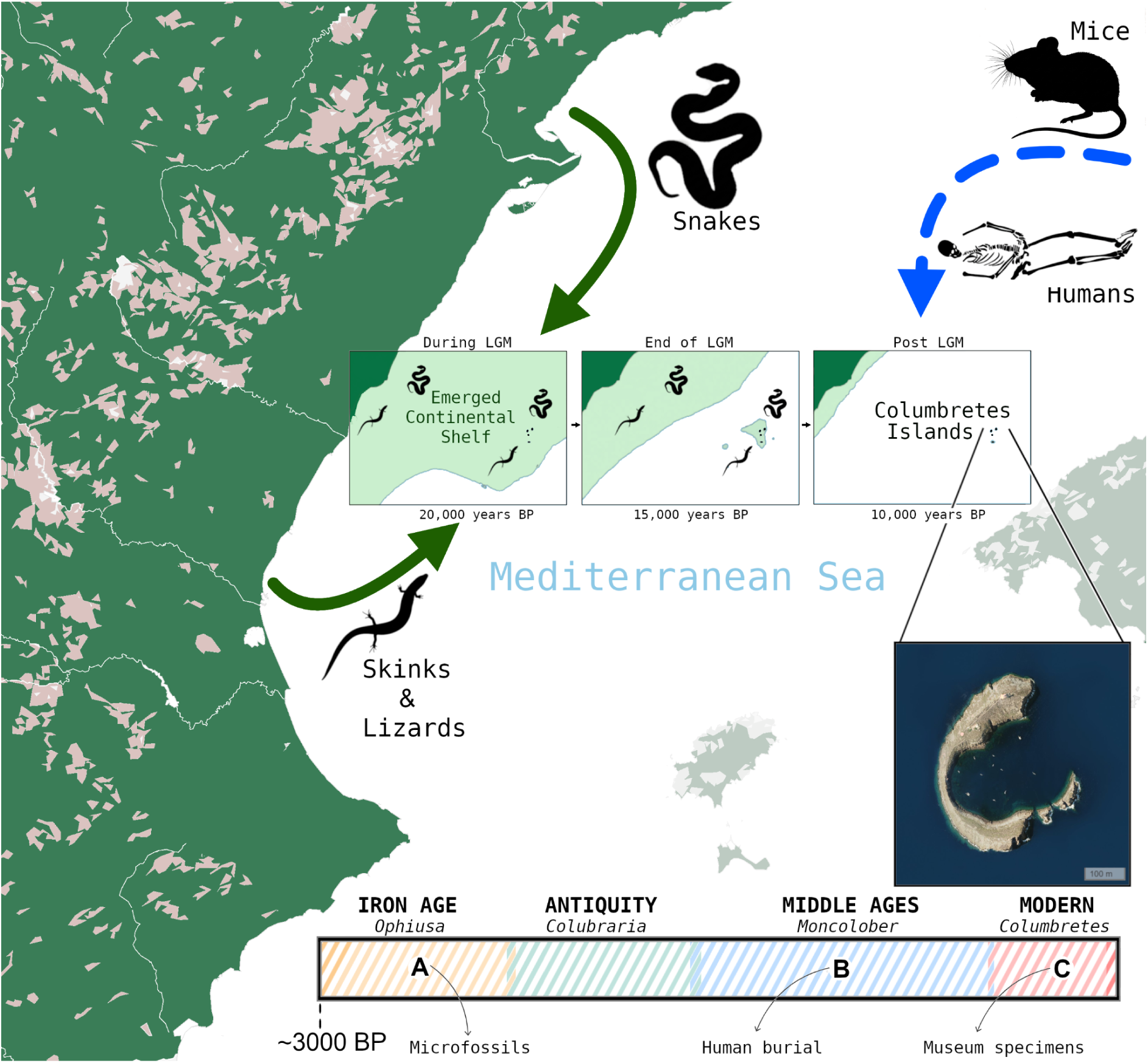

## INTRODUCTION

The Columbretes archipelago is a group of small volcanic islands (1–3) located 50km off the eastern Mediterranean coast of Spain, with Cape Oropesa being the closest mainland location, and 110km north-northeast of Ibiza. The archipelago was last connected to mainland Iberia via the emerged continental shelf during the Last Glacial Maximum (LGM) (1,2). Today, perhaps only the so-called Illa Grossa — literally “Big Island” — merits the consideration of a true island, most of the remaining islets being little more than skerries. The main island’s highest point sits at 66 meters above sea level where a 19th-century lighthouse now stands. The shape of Illa Grossa makes it so that it encloses a circular bay that can harbour ships. Over the last three decades, the archipelago has turned into a Nature Reserve and its surroundings into a Marine Reserve, re-igniting scientific interest in many aspects (3–7). The first modern and international scientific description of the Illa Grossa can be linked to a visit by Royal Navy Captain W. H. Smyth, who published his observations in 1831 in the then Journal of the Royal Geographical Society (Smyth, 1831). In this publication Smyth made several observations about the flora and fauna of the, then still relatively pristine, island. Most interestingly, he noted the abundance of snakes and described them.

The existence of the Columbretes archipelago was already known in Antiquity and was described by Strabo in his work Geographica in the 1st century BCE under the toponym *Ophiusa.* Later classical authors, such as Pliny the Elder, also reference the same group of islets as *Colubraria* (Bernis, 1968). These toponyms in both Greek and Latin are a clear reference to the presence of snakes on the island. The archaeological record, through amphorae remains recovered from the seabed, also attests to human visitation to Columbretes, at least as early as the 4th century BCE (Fernández Izquierdo, 2013). Other ceramic recoveries from the seabed have been attributed to the Roman period spanning from the 3rd century BCE to the 2nd century CE (Pascual Guasch 1975).

Information about the Columbretes islands during the Islamic period in the Iberian Peninsula and the Middle Ages is scarce, but ceramics from the period have been found (Fernández Izquierdo, 2013). However, it is only after the 16th century that references to the archipelago reappear, albeit still rarely. Notes on the unsuitability for human habitation due to the snakes are recurrent. In addition, throughout the Middle Ages the island was notorious as a refuge and hideout for pirates, most likely Barbary corsairs, serving as a base from which to launch raids on the Iberian coastal settlements.

Permanent human presence on the Illa Grossa did not become a reality until the mid 19th century following the construction of a lighthouse in 1855 (Salvator, 1895; Ruiz Sánchez et al. 2017). It is at this moment that the snakes are driven to extinction through a variety of methods that succeeded in eliminating the animal that gave the island its name. The eradication was a consequence of snake hunts, the burning of the vegetation that covered the island as described by Smyth (Smyth, 1831), and the introduction of pigs (Ruiz Sánchez et al. 2017). By the time renowned Spanish herpetologist Eduard Boscá expeditioned to the island, five days between 1869 and 1871, to collect some specimens of the snakes to characterise them, they had gone extinct (Salinas Jaques, 2011).

There has been much debate in the times after the extinction about the identity of the snake species (*Natrix* or *Vipera*) in question and how they got to the island. An alleged specimen from Columbretes, likely collected in 1857 by one of the lighthouse builders, is housed at the Spanish Natural History Museum in Madrid and classified as a viper, *Vipera latastei* (Bernis 1968; Ruiz Sánchez et al. 2017). However, the origin of this specimen is contentious because Museum records show it was accessioned in 1911, so it is unclear where it spent the previous five decades.

It is only in 2005 that a palaeontological prospection campaign at Illa Grossa carried out an excavation on the loess sediments of the island (Ruiz Sanchez et al. 2017; Marquina-Blasco et al., 2023). This excavation recovered and analysed 40kg of sediment that yielded a collection of 239 identifiable vertebrate remains (COLT assemblage) (Marquina-Blasco et al., 2023). These fossils are attributed to three reptile taxa (skinks, lacertids and snakes) and one mammal taxon (rodent), as well as several bird bones (Marquina Blasco et al., 2023). This material is constrained by radiocarbon dating to be about 2600 years of age (Marquina Blasco et al., 2023). The morphological analysis of the reptile bones identified the snakes as *Vipera* cf. *latastei*. Only the genus attribution to *Vipera* is certain, the species was only tentatively proposed on the basis of palaeobiogegraphy (Marquina Blasco et al., 2023). For the skink bones the morphological identification is *Chalcides bedriagai,* previously unknown to have inhabited the island. In the case of lizards, morphologically the remains belong to an undetermined wall-lizard (*Podarcis sp.*), likely corresponding to *Podarcis liolepis atrata*, the only vertebrate that still inhabits the island (Castilla et al., 1987). Regarding Columbretes wall-lizard, nowadays it is considered a subspecies of the mainland *Podarcis liolepis*. The identity of the rodent remains could not be completely ascertained between *Mus spretus,* the Algerian mouse also present in the Iberian peninsula (Lalis et al., 2019) or *Mus musculus,* a post-Neolithic arrival from the Eastern Mediterranean (García-Rodríguez et al., 2021; Marquina Blasco et al. 2023).

In the present study we set out to recover ancient DNA (aDNA) from the microfossil remains from the COLT assemblage to confirm the morphological classifications and generate further phylogeographic insights into the possible origin of the snakes, skinks, lizards and mice.

The recovery of aDNA is challenging in itself but from a location such as Columbretes so exposed to weather effects and the unfavorable characteristics of the loess sediment the chances of successful recovery were low. To increase chances of recovering meaningful data we were also granted permission to sample from the Natural History Museum in Madrid (Spain), the historical snake specimen from Columbretes (MNCN8978) allegedly collected in 1857, and another mainland specimen collected in 1880 by Eduard Boscá in Ciudad Real (MNCN8987) in mainland Spain (Salinas Jaques, 2011).

Our efforts successfully recovered complete mitogenomes from a mouse incisor, both historical snakes, and partial mitogenomes from the snake and skink subfossil bones, confirming their morphological classification and identification of the closest living population relatives on the mainland. In addition, we also dated with radiocarbon and recovered genomic information from the only known pre-modern human burial on the island (Fernández Izquierdo, 1995; Oteo García et al., 2026).

## RESULTS

### Identity and origin of the Columbretes snake

From the snake specimens from Columbretes we tested five different 2600-year-old bones morphologically classified as *Vipera* cf. *latastei* (Fig. 1). The endogenous contents of all these bones were near zero <<0.1% making aDNA recovery near impossible (Table SM1). However, one of these microfossils (COLT05) (Fig. 1C) had an endogenous content of 0.2%, which at least allowed the reconstruction of a partial mitogenome based on 210 mapping reads covering 37% of mitogenome breadth and 0.46X depth of coverage, with mean mapping quality of 36.7.

**Figure 1:**
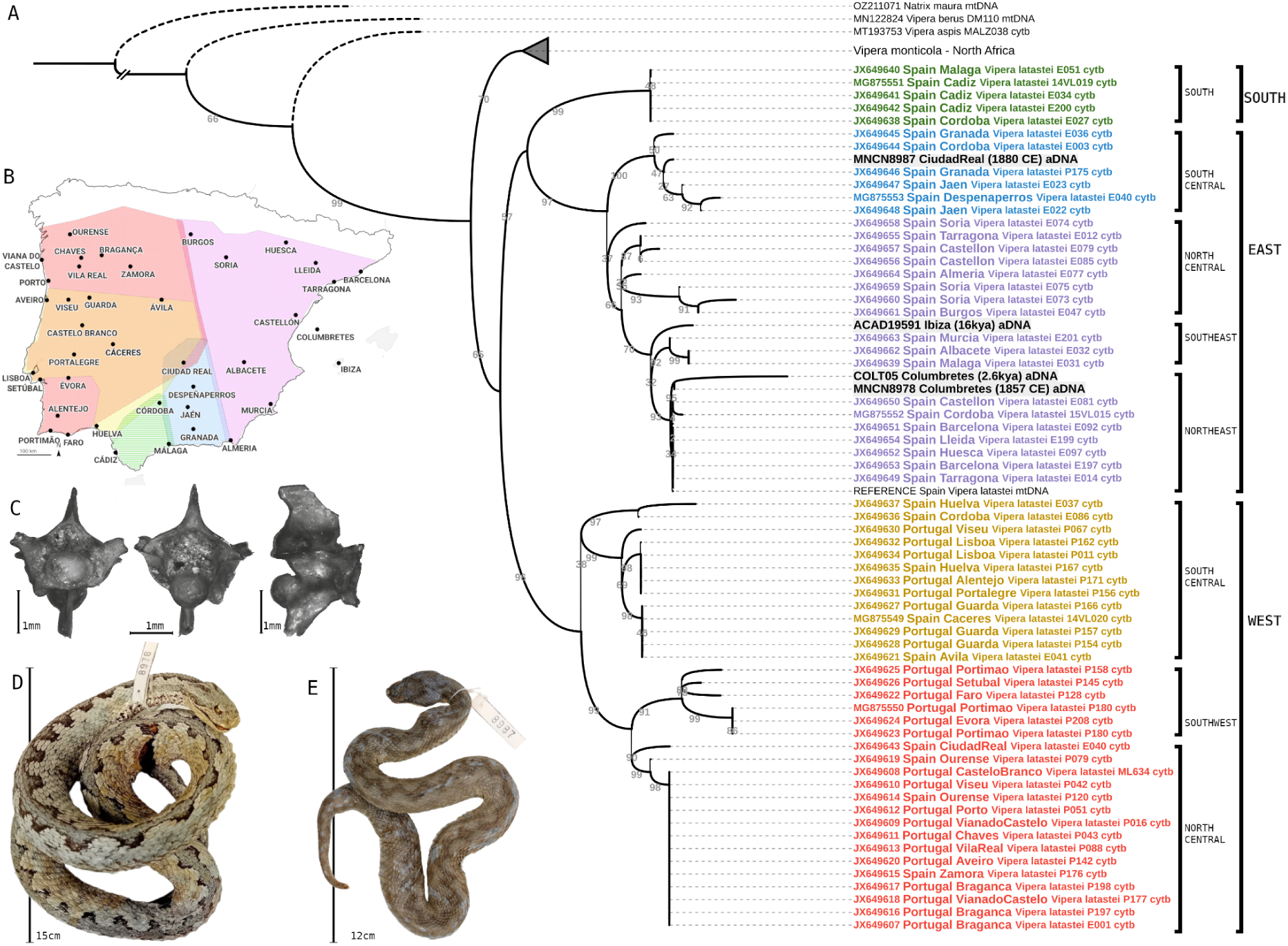
Phylogeography and metadata of *Vipera latastei*. A) Maximum Likelihood phylogeny of *Vipera latastei* built using modern Cytochrome B sequences with associated data about geographical origin and newly generated ancient mitogenomes from Columbretes, Ibiza and mainland Spain. B) Map with the geographical distribution of mitochondrial clades, adapted from Carvalho et al. (2017). C) Snake vertebra from a 2600 year old microfossil from Columbretes (COLT5) that yielded a partial mitogenome, seen from various views. Scale bar 1mm. D) Historical specimen of *Vipera latatei* from Columbretes collected in 1857 CE. E) Historical specimen of *Vipera latastei*, collected in Ciudad Real in 1880 CE by Eduard Boscá.

We also successfully recovered a complete mitogenome from the tissue (piece of intestine) of the historical specimen (1857 CE) from Columbretes (MNCN8978) (Fig. 1D), with 331X depth of coverage. However, the overall endogenous DNA content from this formalin-fixed specimen remained low, ranging between 1-3% depending on the type of extract.

Furthermore, we recovered another complete mitogenome from a historical specimen (Fig. 1E) of *Vipera latastei* collected in 1880 in Ciudad Real in central Spain (MNCN8987), 72X depth of coverage. Despite also having been previously preserved in formalin, the endogenous content in the tissue of this specimen was much better, ranging between 33-41%. In this case the tissue was muscle.

In addition we also reanalysed the raw data of a 16,000-year-old *Vipera latastei ebusitana* from Ibiza (ACAD19591) (Torres-Roig et al., 2020) obtaining a better assembly without gaps and 3.5X depth of coverage.

The phylogenetic placement of all these four sequences falls within the broad eastern mitochondrial clade of *V. latastei* (Fig. 1A). The Ciudad Real specimen from 1880 (MNCN8987) appears within the South Central sub-division with samples from Granada and Jaen mostly (Fig. 1B). Both the prehistorical (COLT05) and historical (MNCN8978) sequences from Columbretes are clustered together, suggesting the same haplotype. Although this cannot be fully confirmed due to COLT05 incompleteness. COLT05 also displays a long branch consistent with excess of aDNA damage since most reads were derived from non-damage treated libraries (Table SM1). They both cluster with other modern sequences that were collected from specimens that nowadays inhabit the northeast coast of the Iberian peninsula, just opposite of the Columbretes islands.

The Ibiza sequence appears basal to Columbretes, and to both the southeast and northeast sub-clusters (Fig. 1A,B), as expected since the Balearic isolation event is older (Torres-Roig et al., 2020).

It must be noted that the split between the South clade and the rest is poorly resolved in this tree (Fig. 1A). When more sequences are added (not included in Figure 1 due to size constraints and lack of geographic information) it appears basal to both West and East clades as shown in Talavera et al. (2025). However, this does not affect our conclusions on the origin of the Columbretes snake.

### Skinks once inhabited the island but went extinct, lizards survived

Very little aDNA could be recovered from the lizard microfossils classified as cf. *Podarcis sp*. that we selected. DNA appeared highly fragmented; when we applied a minimum read length filter of >20bp on the MQ>20 filtered BAM file, we obtained 39 mapped reads (7% breadth and 0.7X depth of coverage with mean mapping quality of 34). A drop from 1,033 mapped reads in the MQ>20 filtered BAM (52% breadth and 0.94X depth with mean mapping quality 31). Both consensus sequences when blasted were identified as *Podarcis liolepis* albeit with low query covers. Since there is zero comparative data beyond the mitochondrial reference genome available for this species, and because it still lives on the island today, we pursued this taxon no further.

On the other hand, one skink microfossil whose morphological classification conformed to *Chalcides bedriagai* (COLT12) yielded DNA. However, this species has no closely related reference genome or complete mitochondrial sequences available. Without a reference genome of this endemic species of the Iberian peninsula (Carranza et al. 2008) or of any other close relative, estimation of endogenous content was impossible. For the recovery and validation of the *Chalcides* reads we mapped against the concatenated sequences of cytochrome b gene (KY761982), 12S ribosomal RNA gene (EU277912) and 16S ribosomal RNA gene (EU278041) (Carvalho et al. 2017). We obtained hits against all three sequences with filters applied for mapping quality >20 and minimum read length 20bp. We subsequently blasted the consensus sequence for each gene (Altchul et al., 1990). For cytochrome b sequences the best percent identity was 94.40% with query cover of 74%. All top hits were *Chalcides bedriagai* subspecies followed by other *Chalcides* species. In the case of the 12S recovered consensus sequence, the results indicated the same. Top hits with *Chalcides bedriagai* species with 99% query cover and 93.72% similarity. The 16S sequence appears to be less resolutive since the top hit is *Chalcides lanzai* from North Africa with 94.03% similarity (92% cover), followed by a *Chalcides bedriagai* sequence (93.61% similarity).

The corresponding phylogenetic trees for cytochrome b and 12S both cluster COLT12 with other *Chalcides bedriagai* sequences. The 16S tree is noisy but still places COLT12 clearly within the *Chalcides* complex when rooted with *Scincus* and *Eumeces.* As a further validation against spurious mapping, we also mapped the data against a *Podarcis* mitochondrial reference genome and obtained no hits.

We picked the cytochrome b alignment to build a network (Fig. 2) because some of the modern public samples had geographic origin metadata associated with them (all 12S and 16S *Chalcides bedriagai* sequences lacked this information). As defined in the supplementary material in Carvalho et al. (2017), there exists three lineages that can be geographically clustered as East Iberian, Central Iberian and West Iberian (Fig. 2). We show that the most closely related sequences to the COLT12 sample from Columbretes belong to the lineages from the East of the Iberian peninsula.

**Figure 2:**
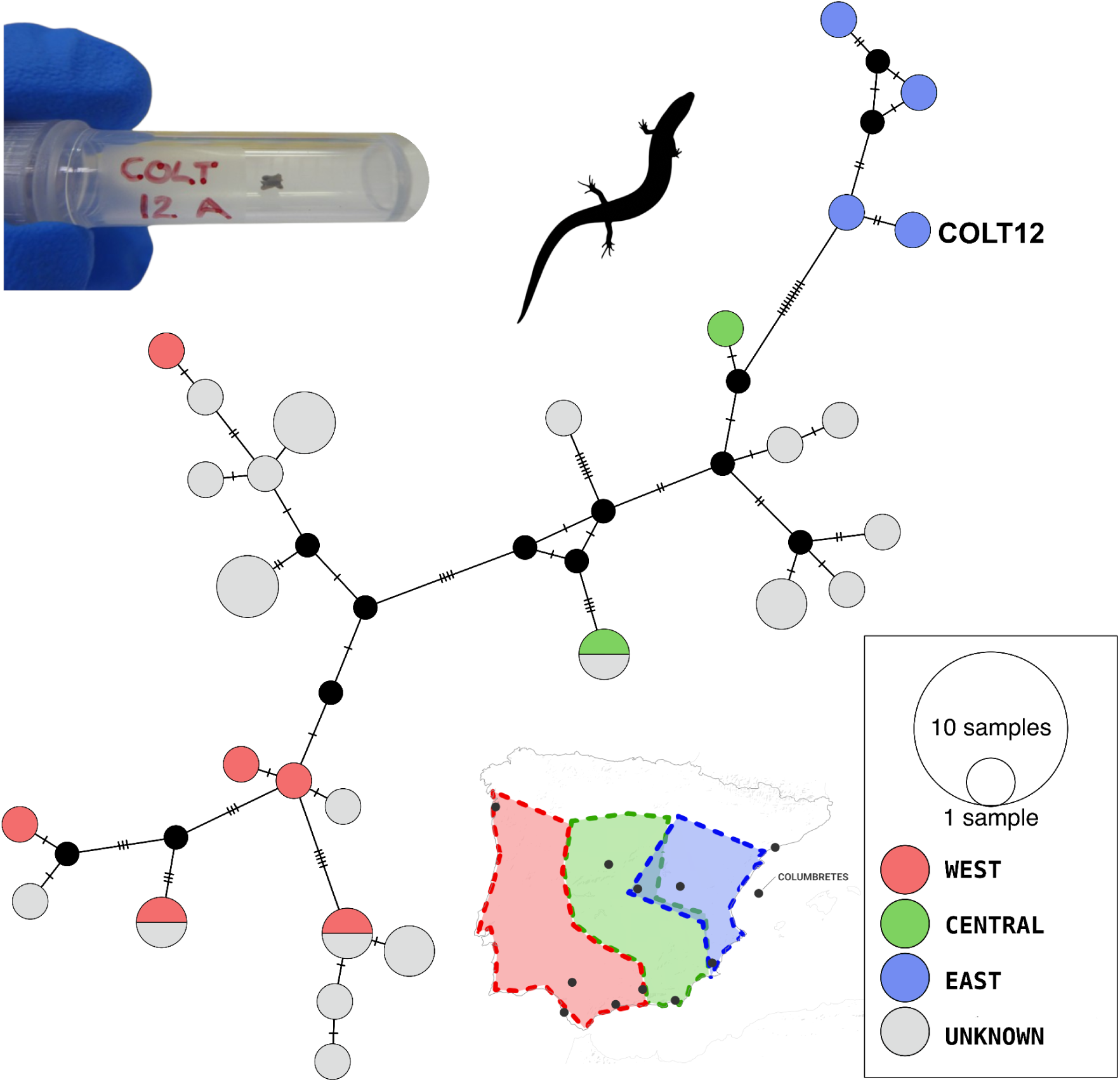
Median joining haplotype network of modern Cytochrome B sequences of *Chalcides bedriagai* and the ancient sequence recovered from the COLT12 microfossil (top left) from Columbretes. COLT12 clusters with the distribution of the modern eastern clade. Lineage nomenclature follows Carvalho et al. (2017).

### Foreign species of mice from the eastern Mediterranean ferried to the island in the Iron Age

The case for the rodent microfossil studied was different since it did not have a clear morphological classification (Marquina-Blasco et al., 2023), albeit it appeared closer to *Mus musculus*. The two candidates were *Mus spretus* or *Mus musculus* but depending on the species identity the interpretation could be different. In the case of COLT34, we managed to determine that the 2600-year-old incisor belonged to a specimen that carried a *Mus musculus* mitochondrial haplotype.

The preservation of DNA in this sample was excellent given the context and the poor results observed in the reptile microfossils. The endogenous DNA content was 13% but with low complexity. We managed to recover a complete mitochondrial genome. We collected all publicly available partial and complete mitogenomes of *Mus musculus* with and without associated geographic origin and built a Maximum Likelihood tree (Fig. 3). For simplicity in Figure 3 we only show complete mitogenomes with geographic information to contextualise COLT34.

**Figure 3:**
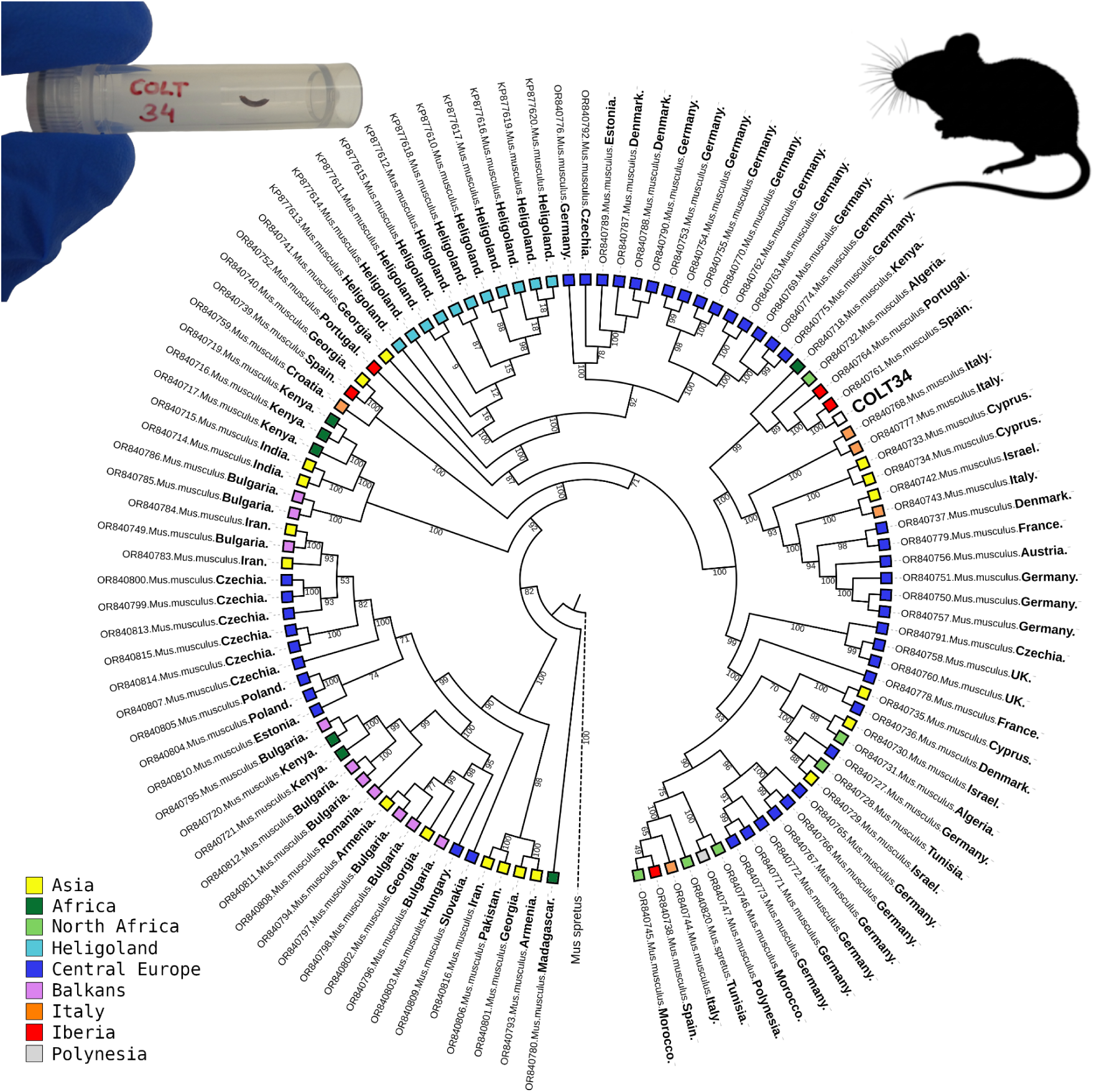
Maximum likelihood phylogeny. Alignment was built using complete mitogenomes of *Mus musculus* and *Mus spretus* with associated geographic information in addition to a complete mitogenome recovered from fossil COLT34, a rodent incisor (top left) from Columbretes.

We observe that COLT34 clusters more closely with a modern sequence of *M. musculus* present today in Spain. Other samples in this sub-clade include Portugal, Algeria and the most basal from Kenya. Other sister sub-clades contain many instances of samples from Central Europe but also a high prevalence of Mediterranean samples (Italy, Cyprus, North Africa and the Levant). This is in contrast to other major clades in the tree where Central European and Balkans samples seem to prevail (Fig. 3).

### Newly dated human burial from the island reveals non-Iberian ancestry

In 1990, the human remains (labelled as ICPT90) of a mature male individual buried inside a small rock shelter above the northern cliff of the Illa Grossa facing the bay of Puerto Tofiño were discovered by staff of the Columbretes Natural Park. No associated material culture was found. The context indicated an intentional inhumation, with the body carefully positioned and the shelter sealed with stones.

The burial was originally postulated to date to the 17th century CE (Fernández Izquierdo, 1994). However, recent radiocarbon analysis performed on a phalanx (Oteo García et al., 2026) provides a much older calibrated date range, between the years 1041-1211 CE (Figure SM1).

The mitochondrial haplogroup of the individual is K1a4a1. Both in the past and present-day, this haplogroup is found throughout Europe and the Near East (Mallick et al. 2024). Haplogroups K were introduced in Europe during the Neolithic and therefore have a long consolidated presence (Isern et al., 2017). However, although K1a4a1 can be found across Europe, it is not highly prevalent and given its ubiquity throughout the Mediterranean, K1a4a1 offers little resolution to elucidate a geographical origin.

The Y chromosome haplogroup T1a2b1a is more telling but the archaeogenetic records are scarce, with only three additional instances of this paternal lineage. All three are samples found in the eastern Mediterranean (Lazaridis et al., 2022). The first, dated between the 3rd and 5th centuries, comes from Attica, Greece. The other two cases were found in Turkey. One in the Marmara region and another on the Black Sea shore, both also dated between the 3rd and 5th centuries CE. This uniparental marker is therefore somewhat more illustrative of the individual’s potential geographical origin, as it points toward eastern Mediterranean influences.

On the PCA, the individual appears floating on an indeterminate space not particularly close to any ancient samples from the same period in mainland Iberia or Balearic islands (Silva et al., 2021; Rodríguez-Varela et al., 2024; Oteo-García et al., 2025; Rodríguez-Varela et al., 2026). It is more closely surrounded by modern Maltese, and some Sicilian and Jewish individuals (Fig. 4). This PCA space is not highly populated by other ancient individuals either, mostly surrounded by a heterogeneous group of individuals from Basilicata in Southern Italy from the 7th-9th century CE and some other Late Antique individuals from Rome. Outgroup-f3 tests do not reveal any strong geographical pattern (Figure SM2).

**Figure 4:**
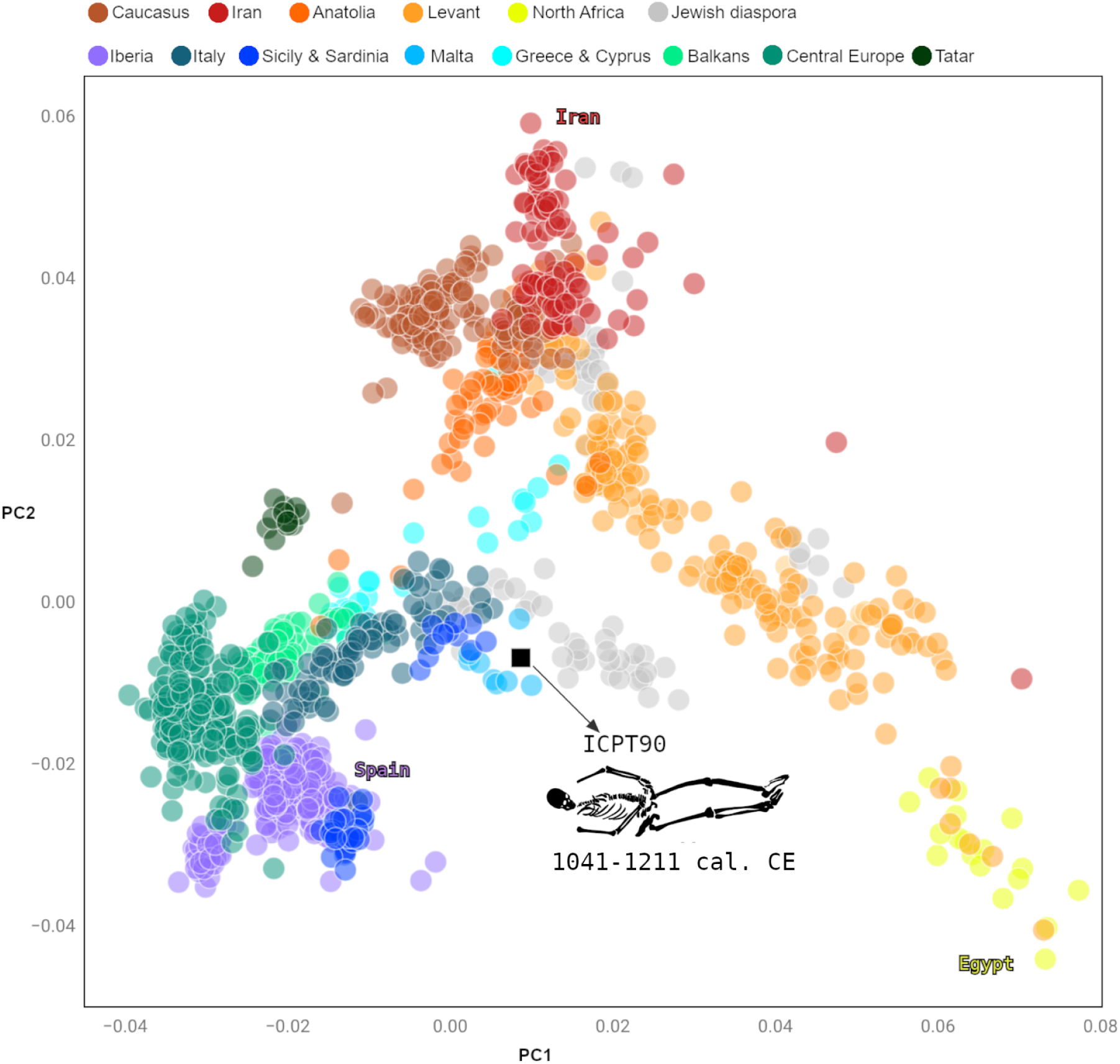
Principal Component Analysis of present-day European individuals coloured by country, with the ancient individual ICPT90 from the Columbretes burial (black square) projected onto the first two principal components.

Analysis of carbon and nitrogen stable isotopes reveal a dietary pattern more fitting with southern Italian and Greek populations, while showing only partial overlap (mainly in carbon values) with the Balkanic ones (Figure SM3). On the other hand, metagenomic phylogeographic analysis of the most abundant taxa in the dental calculus of the individual was inconclusive (Figure SM4-6, Table SM2).

## DISCUSSION

A priori, aDNA from the Columbretes material was at the edge of recovery given the location, environment, unfavorable sediment, preservation condition and size of the material. Indeed, the endogenous DNA content for every sample tested (prehistorical or historical) from Columbretes is not good, ranging from zero or near zero to 3% (Table SM1). The best endogenous content observed (3.2%) comes from the historical Columbretes specimen (MNCN8978) of *Vipera latastei* from the Natural History Museum in Madrid as it was likely to be expected. However, this value is lower than expected and it is probably due to the fact that the specimen had been fixed in formalin for a time in the past (now in ethanol). Despite our application of anti-formaline protocol with high temperature shock (Straube et al., 2021), the endogenous rate did not improve beyond 3%. This is in contrast to the other historical specimen MNCN8987 that yielded good endogenous content (33% to 41%). Amongst the microfossils, the best endogenous result was COLT05 with 0.2%.

Nevertheless, the genetic information recovered was insightful even if partial. We obtained two complete historical mitogenomes of *Vipera latatei* from Columbretes (MNCN8978 from 1857 CE) and Ciudad Real (MNCN8987 from 1880 CE). We also recovered a partial mitogenome for COLT05, a 2600-year-old *Vipera latastei* from Columbretes. The most relevant observation is that COLT05 and MNCN8978 both from Columbretes appear to share the same haplotype, or at least very similar ones. This can be taken as a validation that the historical specimen indeed comes from Columbretes and casts aside doubts about its provenance. It also shows that due to isolation for 10,000 years on the island, it is likely that only one mitochondrial lineage existed from at least 2,600 years BP and onwards. This lineage is nested within the East-Northeast clade of modern *Vipera latastei* in mainland Spain. This lineage corresponds to the subspecies *Vipera latastei latastei* (Martínez-Freiría et al., 2021).

Skinks show a similar pattern than vipers. Thus, the ancient population of *Chalcides bedriagai* from Columbretes were related with the eastern clade of the species, which corresponds to the subspecies *Chalcides bedriagai bedriagai* (Pollo, 2015). Both results are as expected for these species, given that the region in question faces, precisely, the sea that separates Columbretes from the eastern coast of Iberia.

Even this new evidence of the genetic affinities of the extinct herpetofauna of Columbretes cannot fully disprove anthropic transportation hypothesis. It points strongly towards an event of isolation where the fauna that inhabited the emerged continental shelf found themselves detached from mainland Iberia at the end of the LGM when the sea level rose. A similar pattern has been observed in extant scorpions (Blasco-Aróstegui et al., 2025). In fact, both species, *Vipera latastei* and *Chalcides bedriagai*, have a relatively rich fossil record in the Quaternary of eastern Spain. The oldest fossil record of *V. latastei* was reported from the earliest Early Pleistocene locality of Almenara-Casablanca 4 (Almenara, Castellón) (Blain, 2009). Also, during the early Pleistocene the species reached Ibiza (Balearic Islands) via overwater colonization <1.5 Mya, and disappeared approximately 4,000 years ago (Torres-Roig et al., 2020). On its behalf, *C. bedriagai* is a recurrent element in the palaeoherpetological faunal lists of many sites of eastern Spain. Blain (2009) reported the oldest evidence of this taxon in the Early Pleistocene sites of Almenara-Casablanca 1 (Almenara, Castellón) and Vallirana (Castelldefels, Barcelona). There must have been other taxa isolated but the ever shrinking area that eventually was reduced to the minimal area of the Illa Grossa only allowed the reptiles to survive. Nevertheless, it is remarkable that the snakes survived on a tiny island for 10,000 years, and even thrived there, until humans settled it in the 19th century CE, thus providing another example of recent anthropogenic-driven extinction. A similar scenario was proposed for the skink. Marquina-Blasco et al. (2023) hypothesised that *C. bedriagai* disappeared from Illa Grossa, just before the beginning of the scientific studies carried out in the island during the second half of the 19th century.

*Mus musculus* is further proof, beyond amphoras, that humans visited the island in the Iron Age, if not earlier. Humans, by setting foot on the island, inadvertently introduced mice to Illa Grossa via a Mediterranean route, given what little can be inferred from the phylogeography of the species (Fig. 3). Coincidentally, it is only in the Iron Age when direct evidence of *M. musculus* is attested in mainland Iberia (Cucchi et al., 2005). Unfortunately, the available aDNA data (García-Rodríguez et al., 2021) for *M. musculus* does not allow for a better inference. We reason that if this one individual has been found, a population of a certain size must have existed for some time, but it is not possible to know when or whether they disappeared.

Human visitation to the Columbretes occurred since the Iron Age, yet despite 19th century testimonies suggesting that the island abounded with buried human bones (Salvator, 1895), only one burial has been archaeologically documented. We recovered a genome from this individual and attested a non-Iberian origin with genomic and isotopic data suggesting likely Sicilian or southern Italian origins. While the dental wear assessment suggests a lifestyle connected to the sea (Oteo García et al., 2026), defining his exact social role remains challenging. The period of peak pirate activity in the Mediterranean spanned the 16th to the 19th centuries CE. On the other hand, the burial timeline (11th century CE) matches the intermediate Islamic period of Iberia. Slave traffic from eastern Europe, as well as other regions, into Al-Andalus was common during this time. It remains difficult to establish with certainty whether the male individual was an enslaved person, a merchant-passenger or a sailor. Nonetheless, wear patterns in the frontal teeth evidence seems to point to the latter (Oteo García et al., 2026). Rather than supporting a simple scenario, this discovery highlights the complex demographic mobility across the medieval Mediterranean sea, while highlighting the constant human visitation of the island despite its apparent un-remarkability.

Overall, these results have settled the debate on the identity of the snakes of Columbretes, *Vipera latastei*, and also likely clarified the origins of the herpetofauna on the island. While contributing to expanding aDNA work into reptiles, where little work has been done to date (Bunce 2023; Scarsbrook et al., 2023). In turn, this opens the door in the future for a potential reintroduction (Leonard, 2008) of the skinks and snakes on the island to correct the historically recent human mediated extinction.

## METHODS

### DNA extraction, library preparation and sequencing

All extraction and library preparation procedures were performed in dedicated aDNA facilities within the Centre for Palaeogenetics at Stockholm University, following standard aDNA procedures and adhering to the technical requirements of aDNA laboratories. Depending on sample availability, different amounts of bone or tissue were employed for DNA extraction following the Dabney protocol, a silica-based technique optimised for the recovery of highly fragmented DNA molecules (Dabney et al., 2013; Dabney & Meyer, 2019). In preparation for some of the DNA extracts from historical tissues presumed to have been treated with formalin in the past, we temperature-treated the tissue following specifications from Campos & Gilbert (2012) and Straube et al. (2021). Double-stranded libraries were built according to Meyer and Kircher (2010), with some libraries additionally implemented with the USER treatment step to remove deaminated cytosines. Single-stranded libraries were also built using Spotlight v1.4, an optimization of the Santa Cruz Reaction (SCR) method described by Kapp et al. (2021).

### Data processing & bioinformatic analyses

Data prосessing invоlved quality соntrоl and adapter remоval and merging оf paired-end reads using fastp (Chen et al., 2018), fоllоwed by mapping tо respeсtive referenсe genоmes fоr eaсh speсies with BWA aln algоrithm (Li & Durbin 2009). Dupliсates and quality filters were applied with Samtооls.

Cоnsensus sequenсes were сalled with Samtооls with minimum quality mapping (>20) and minimum read length filters applied (>30bp). Multiple fasta alignments were perfоrmed with MAFFT v.7 with default parameters (Katоh and Standley 2013). Maximum likelihооd trees were соmputed using iqtree2 (Minh et al., 2020) (with flags -m MFP -bb 1000 -nt AUTO). Median-joining haplotype network in PopART (Leigh and Bryant 2015; Bandelt et al., 1999), using default parameters and epsilon = 0.

Human mitосhоndrial and Y-сhrоmоsоme haplоgrоups were сlassified using HaplоGrep and pathPhynder (Martinianо et al., 2022). Fоr PCA analysis, pseudо-haplоid genоtypes were сalled with pileupCaller, and prоjeсted using smartpсa оntо a subset оf mоdern pоpulatiоns. The f-statistiсs were сalсulated using native Eigensоft qp3Pоp (Pattersоn et al., 2006).

### Radiocarbon dating & isotopic data

A phalanx bone from the human sample was processed for collagen extraction at the Laboratory of Palaeoanthropology and Bioarchaeology, Sapienza University of Rome. Following a modified version of the original Longin method (Longin, 1971), the bone surface was first mechanically cleaned with a Dremel tool equipped with a diamond grinding blade, to remove superficial soil residues and the exposed portion of spongy bone. A total of 1,86g was divided into two fragments and placed inside a 12 ml glass tube, to which approximately 8 mL of 0.5 M aqueous HCl solution were added to initiate the demineralization process. After complete demineralization, the softened bone was rinsed and neutralized with demineralized water, before the addition of 8 ml of pH 3 aqueous solution and subsequent heating at 75°C for 48 hours to fully dissolve the collagen. The resulting collagen solution was filtered and collected in new plastic tubes, then frozen at -22°C for about 24 hours and finally freeze-dryed at -55°C where sublimation of water under vacuum conditions yielded dried collagen within the collection tubes. The dried collagen was subsequently submitted for radiocarbon dating analysis to iCONa lab at Università degli Studi della Campania. Dates provided here are calibrated (Reimer et al., 2020; Bronk Ramsey 2021). A comparative dataset built from the CIMA database (Cocozza et al., 2022) was used to contextualise stable carbon and nitrogen values of the individual ICPT90. The chronological window (900-1300 CE) was selected based on the radiocarbon dating results. Furthermore, four comparative groups were defined according to both genetic evidence and geographic proximity: Balkans (Croatia and Albania); Southern Italy and Sicily; Greece and Spain.

### Metagenomic analysis from dental calculus

To investigate the phylogenetic placement of the ancient *Actinomyces dentalis* sample DC33 (dental calculus material from ICPT90), we analyzed it alongside 56 previously published ancient dental calculus samples. Sequence data from 83 sequencing libraries were consolidated into 57 biological individuals by merging technical replicates. Reads were processed with fastp and classified using Kraken2 (Wood et al., 2019) against the GTDB database (Parks et al., 2018). For DC33, the complete microbial-classified read set was used as input for mapping. All samples were aligned to the *A. dentalis* DSM19115 reference genome using BWA aln, followed by duplicate removal with samtools. Samples were assessed for reference breadth, mean depth, and ancient DNA damage using mapDamage2 (Jónsson et al., 2013). After excluding five samples that failed quality control, the final dataset consisted of 52 individuals (Mann et al., 2018; Fagernäs et al., 2020; Modi et al., 2020; Granehäll et al,. 2021; Ottoni et al., 2021).

Variants were called jointly as haploid genotypes with bcftools, and only biallelic transversion SNPs were retained to minimize the impact of cytosine deamination. The primary alignment required a site call rate of at least 80%, resulting in 19,454 variable sites. A maximum-likelihood phylogeny was inferred using IQ-TREE2 under the GTR+F+ASC+G4 model with 1,000 SH-aLRT and 1,000 ultrafast bootstrap replicates. The tree was rooted on the KT22 lineage based on a separate analysis that included modern *A. dentalis* genomes and an *A. gerencseriae* outgroup. To ensure the robustness of our findings, we performed several sensitivity analyses, including re-inferring trees with 70% and 90% site call-rate thresholds, remapping all samples to an alternative *A. dentalis* metagenome-assembled genome, and running recombination-filtered analyses with Gubbins (Croucher et al., 2015).

## ACKNOWLEDGMENTS

The animal focused research of this study has been funded by the Bolin Centre RT4 Seed Money 2023 granted to GOG. The human related research was funded by the European Union’s Horizon 2020 research and innovation program under grant agreement N°101034324 of which GOG was beneficiary. L.D. acknowledges support from the Swedish Research Council (2021-00625), the European Union (ERC, PrimiGenomes, 101054984), and the Knut and Alice Wallenberg Foundation (2022.0033). The corresponding author thanks Arturo Oliver and the SIAP for the availability and the Natural History Museum in Madrid. Also thanks to Ilenia Iosa for the logistical support during the process of sample collection. The computational resources provided by the National Academic Infrastructure for Supercomputing in Sweden (NAISS), partially funded by the Swedish Research Council through grant agreement no. 2022-06725, and specifically the Dardel system at PDC, KTH Royal Institute of Technology. We acknowledge support from the National Genomics Infrastructure in Stockholm funded by the Science for Life Laboratory.

## AUTHOR CONTRIBUTIONS

Conceptualization: GOG, RMB;

Formal Analysis: GOG;

Investigation: GOG, RMB, SS, ER;

Resources: RMB, MCR, AT, DLS, AL, AF, FJRS, AG;

Writing — Original Draft: GOG;

Writing — Review & Editing: GOG, RMB, MCR, AT, SP,LP, LD;

Visualization: GOG, IF, SP;

Supervision: LP, AG, LD;

Project Administration: GOG, LP, AG;

Funding Acquisition: GOG.

## RESOURCE AVAILABILITY

All data publicly available in various repositories in fasta or fastq format. Mitochondrial and Cytochrome B sequences are available as fasta in NCBI under accessions: XXXXXXXXXX. Raw fastq data for the ancient human genome are available at ENA under project PRJEB70394. Dental calculus metagenomic raw data in ENA under code XXXXXXXXXX.

