## Supplementary Information for "*Hic sunt dracones*: ancient DNA from the extinct palaeofauna of the tiny Mediterranean archipelago of Columbretes and its implications for human visitation"

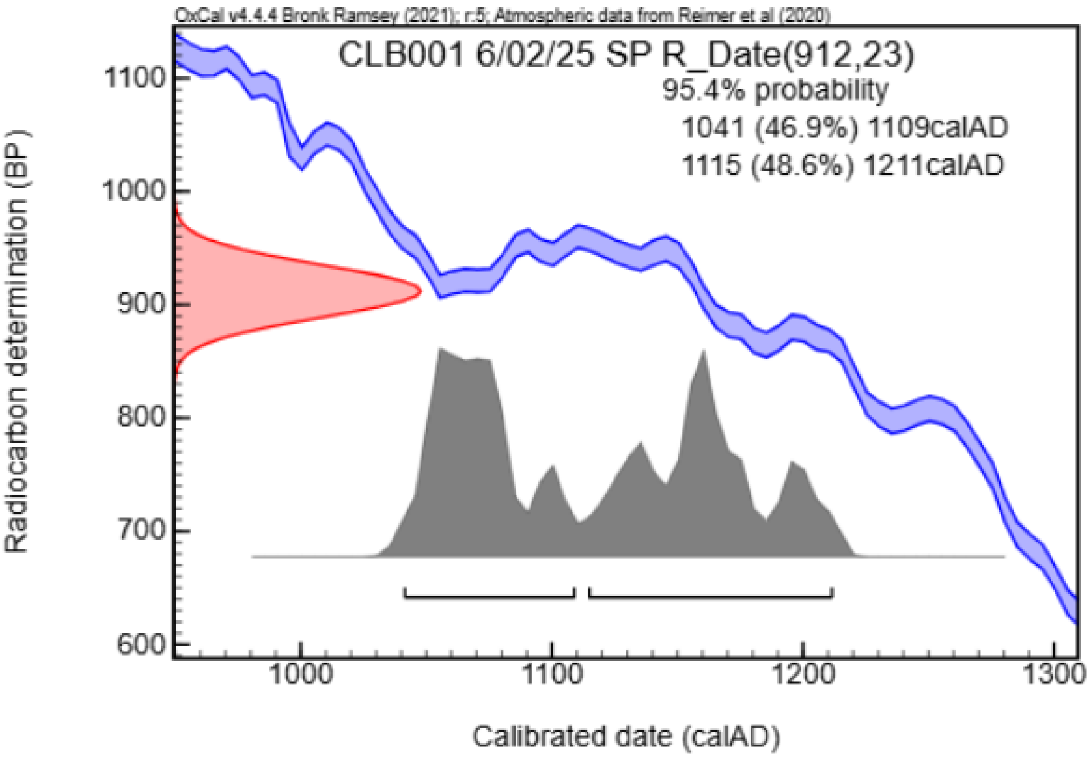


Figure SM8: Radiocarbon date obtained for the human burial in Columbretes (ICPT90-CLB001).

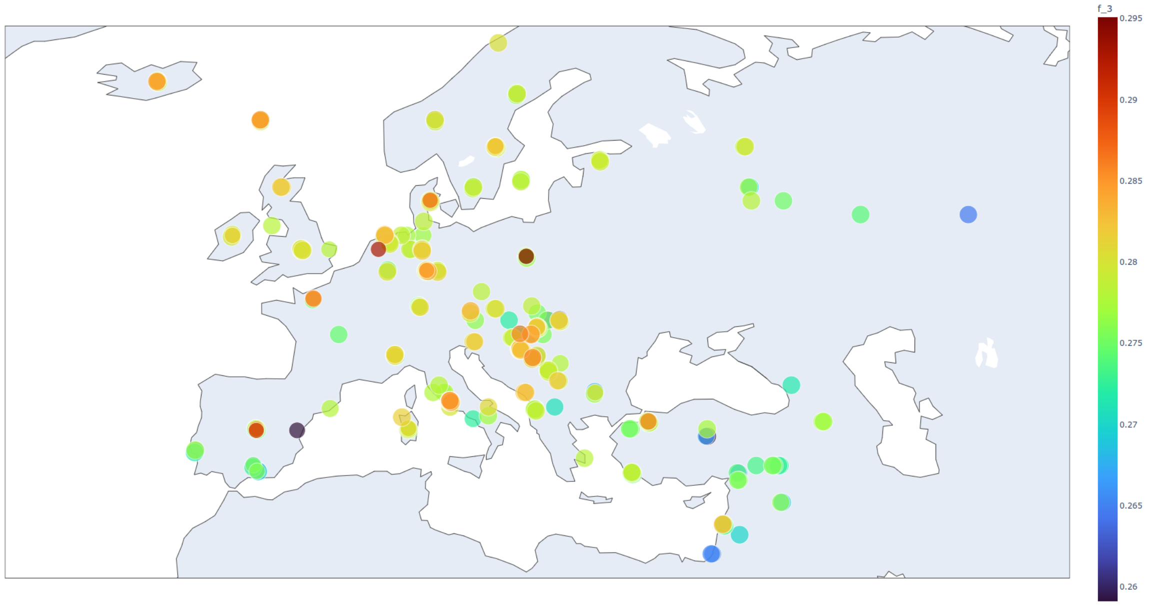


Figure SM2: Outgroup-f3 for ICPT90 with other ancient individuals of similar age and location on map measured as f3(Mbuti; ICPT90, Ind-X).

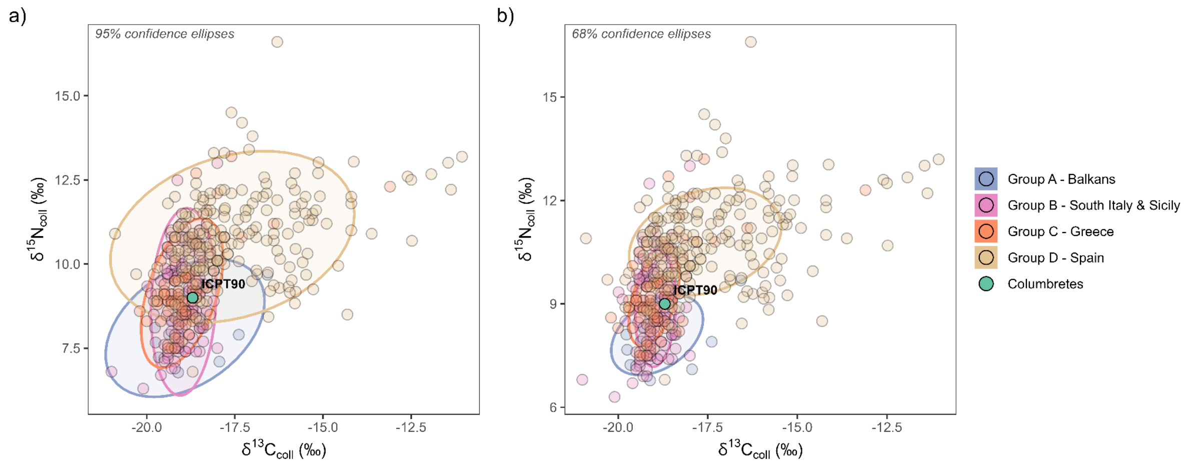


Figure SM3a: d13C and d15N values of the Columbretes individual (ICPT90) compared with regional datasets from the CIMA database for the 900-1300 CE chronological window. Comparative groups selected: A - Balkans, B - Southern Italy and Sicily, C - Greece, D - Spain. At 95% confidence ellipse (a) and 68% confidence interval (b).

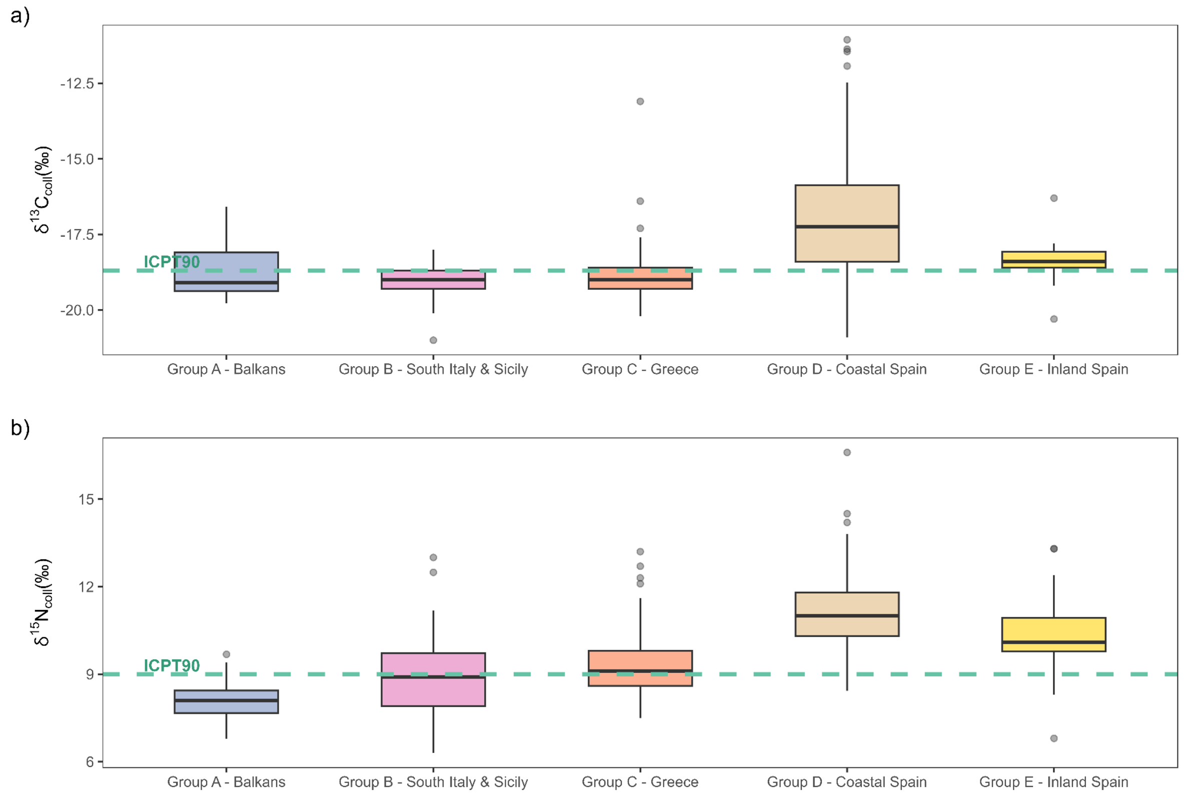


Figure SM3b: Boxplots comparing d13C (a) and d15N (b) values of the Columbretes individual (ICPT90, dashed green line) against regional datasets from the CIMA database for the 900-1300 CE chronological window. Comparative groups selected: A - Balkans, B - Southern Italy and Sicily, C - Greece, D - Coastal Spain, and E - Inland Spain.

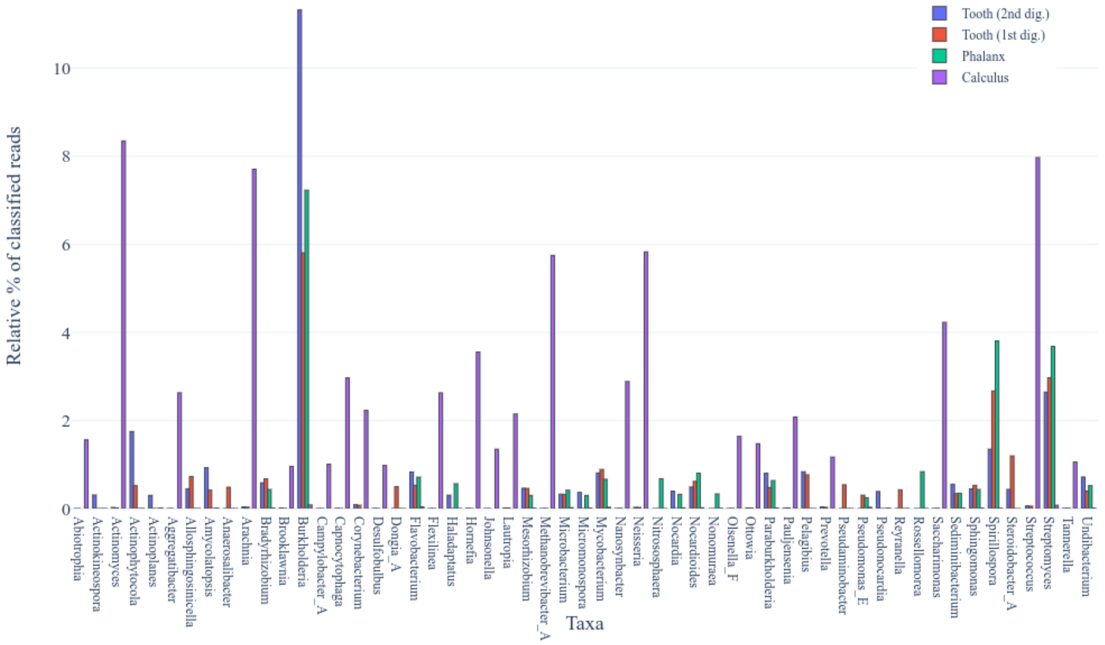


Figure SM4: Metagenomic composition, as classified by Kraken2, of libraries sequenced from various materials (tooth cementum, phalanx and dental calculus) from ICPT90 from where we obtained DNA extracts.

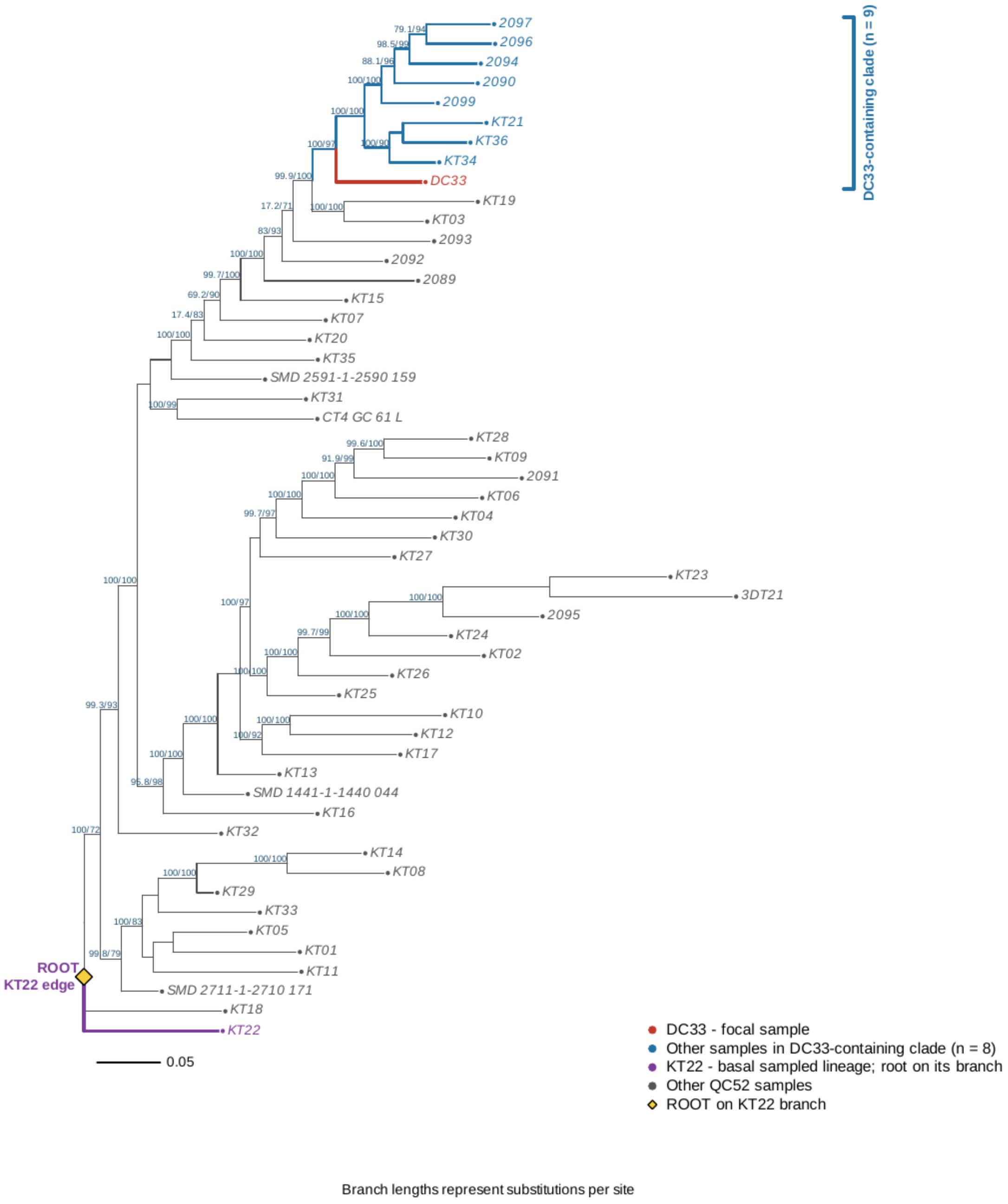


Figure SM5: Phylogeny of *A. oralis* from several ancient samples from Europe of similar age as ICPT90.

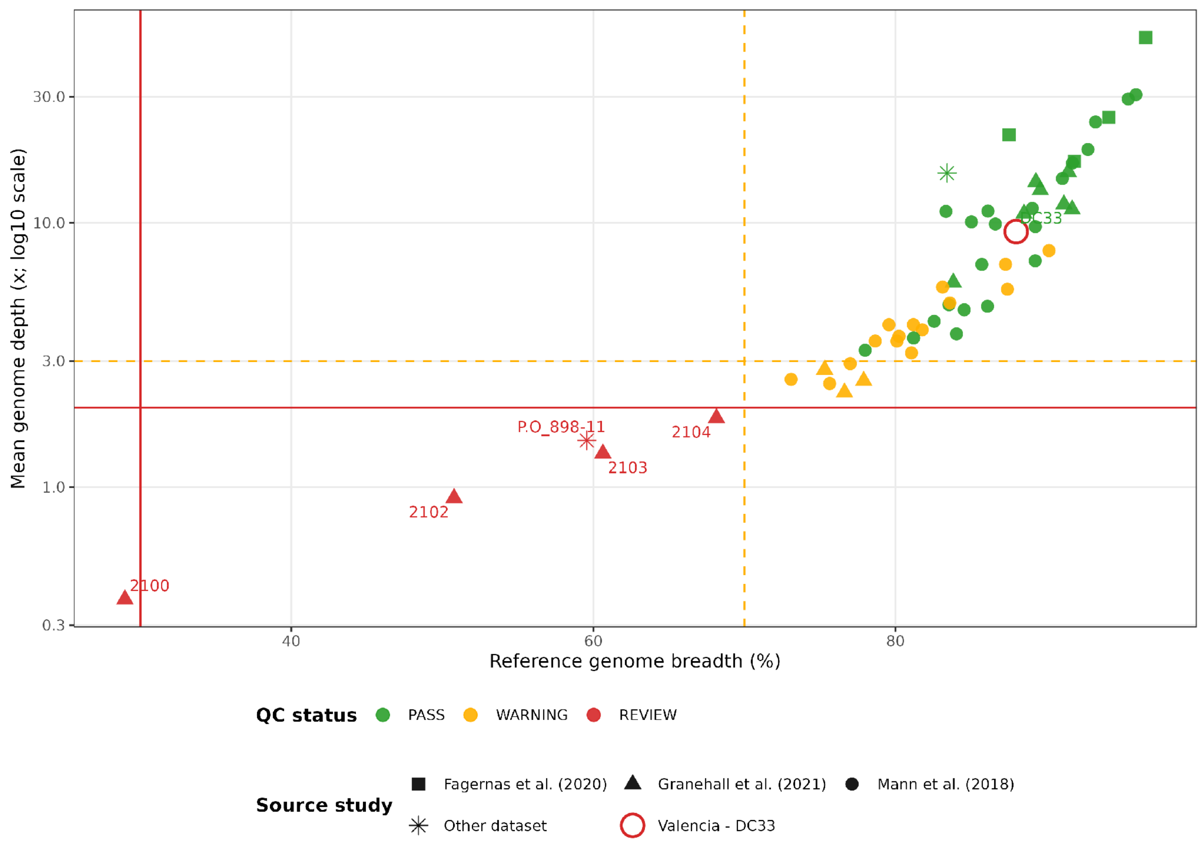


Figure SM6: Sequences quality thresholds across 57 *A. oralis* genomes recovered from various dental calculus samples across various locations. Symbols and IDs in red are excluded from the final results. Dashed lines mark threshold for 70% breadth and 3x depth of coverage.

Table SM1: Endogenous content for all DNA extract from each sample studied here for each animal species.

| Library_ID | Extract_ID | Location | Library_type | Species | Endogenous DNA % |
|---|---|---|---|---|---|
| P34805_1112 | COLT05 | Columbretes | MK non-user | Vipera | 0.271 |
| P35758_1076 | COLT05 | Columbretes | MK user | Vipera | 0.07 |
| P36262_1172 | COLT05 | Columbretes | MK user | Vipera | 0.176 |
| P36262_1173 | COLT05 | Columbretes | MK user | Vipera | 0.208 |
| P36262_1174 | COLT05 | Columbretes | MK user | Vipera | 0.239 |
| P40061_1048 | COLT05 | Columbretes | MK user | Vipera | 0.169 |
| P34805_1113 | COLT06 | Columbretes | MK non-user | Vipera | 0.002 |
| P32015_1092 | COLT10 | Columbretes | SCR | Vipera | 0.002 |
| P32015_1096 | COLT50A | Columbretes | SCR | Vipera | 0.002 |
| P32015_1097 | COLT50A2 | Columbretes | SCR | Vipera | 0.002 |
| P32015_1098 | COLT51A | Columbretes | SCR | Vipera | 0.019 |
| P32015_1099 | COLT51B | Columbretes | SCR | Vipera | 0.001 |
| P32015_1101 | MNCN8978a | Columbretes | SCR | Vipera | 0.143 |
| P33206_1010 | MNCN8978a | Columbretes | MK user | Vipera | 2.242 |
| P36262_1249 | MNCN8978a | Columbretes | MK user | Vipera | 0.995 |
| P32015_1105 | MNCN8978a2 | Columbretes | SCR | Vipera | 0.018 |
| P32015_1102 | MNCN8978b | Columbretes | SCR | Vipera | 0.005 |
| P32015_1103 | MNCN8978c | Columbretes | SCR | Vipera | 0.024 |
| P33206_1011 | MNCN8978c | Columbretes | MK user | Vipera | 2.051 |
| P36262_1175 | MNCN8978c | Columbretes | MK user | Vipera | 1.771 |
| P36262_1250 | MNCN8978c | Columbretes | MK user | Vipera | 1.231 |
| P32015_1104 | MNCN8978c2 | Columbretes | SCR | Vipera | 0.365 |
| P33206_1012 | MNCN8978i | Columbretes | MK user | Vipera | 1.568 |
| P36262_1176 | MNCN8978i | Columbretes | MK user | Vipera | 1.385 |
| P36262_1251 | MNCN8978i | Columbretes | MK user | Vipera | 0.865 |
| P33206_1013 | MNCN8978ii | Columbretes | MK user | Vipera | 3.294 |
| P36262_1177 | MNCN8978ii | Columbretes | MK user | Vipera | 2.895 |
| P36262_1252 | MNCN8978ii | Columbretes | MK user | Vipera | 1.947 |
| P32015_1100 | MNCN8987 | Ciudad Real | SCR | Vipera | 33.096 |
| P35758_1077 | MNCN8987 | Ciudad Real | MK user | Vipera | 38.945 |
| P36262_1178 | MNCN8987 | Ciudad Real | MK user | Vipera | 41.298 |
| P32015_1094 | COLT34 | Columbretes | SCR | Mus | 1.662 |
| P33206_1014 | COLT34 | Columbretes | MK user | Mus | 14.055 |
| P33206_1025 | COLT34 | Columbretes | MK user | Mus | 13.481 |
| P33206_1026 | COLT34 | Columbretes | MK user | Mus | 13.437 |
| P33206_1027 | COLT34 | Columbretes | MK user | Mus | 13.829 |
| P33206_1028 | COLT34 | Columbretes | MK user | Mus | 13.653 |
| P36262_1248 | COLT34 | Columbretes | MK user | Mus | 9.577 |
| P36262_1253 | COLT34 | Columbretes | MK user | Mus | 8.148 |
| P36262_1254 | COLT34 | Columbretes | MK user | Mus | 8.159 |

| | | | | | |
|---|---|---|---|---|---|
| P36262_1255 | COLT34 | Columbretes | MK user | Mus | 8.501 |
| P36262_1256 | COLT34 | Columbretes | MK user | Mus | 9.646 |
| P32015_1093 | COLT12 | Columbretes | SCR | Chalcides | NA |
| P34805_1114 | COLT12 | Columbretes | MK non-user | Chalcides | NA |
| P32015_1095 | COLT48 | Columbretes | SCR | Podarcis | 0.051 |
| P34805_1115 | COLT48 | Columbretes | MK non-user | Podarcis | 0.233 |

Table SM2: Comparative individuals included in dental calculus metagenomic analysis.

| **Samples** | **Location** | **Source dataset** |
|---|---|---|
| KT01-KT36 | Kilteasheen, Ireland | Mann et al. (2018) |
| 2089-2104 | Trentino-Alto Adige, Italy | Granehäll et al. (2021) |
| 3DT21 | York, England | Fagernäs et al. (2020) |
| SMD_* | St. M. de Dulantzi, Spain | Fagernäs et al. (2020) |
| P.O_898-11 | Porticus Octaviae, Italy | Modi et al. (2020) |
| CT4_GC_61_L | Grotta Continenza, Italy | Ottoni et al. (2021) |
